# Prediction from Statistical Learning Aids Auditory Scene Analysis

**DOI:** 10.64898/2026.04.21.719938

**Authors:** Vibha Viswanathan, Srinidhi Narayanan, Ingrid S. Johnsrude, Jenny R. Saffran, Barbara G. Shinn-Cunningham

**Affiliations:** Neuroscience Institute, Carnegie Mellon University, Pittsburgh, PA; Department of Biomedical Engineering, Carnegie Mellon University, Pittsburgh, PA; Department of Psychology, University of Western Ontario, London, ON, Canada; Department of Psychology, University of Wisconsin-Madison, Madison, WI

## Abstract

Statistical regularities support auditory scene analysis across multiple levels. While acoustic regularities like comodulation and harmonicity aid bottom-up perceptual grouping, higher-level regularities like linguistic or musical structure must be learned to form a mental “schema” of statistical patterns. Although learned schemas may benefit comprehension by helping listeners perceptually separate and/or attend to a target sound stream in an acoustic mixture, the underlying mechanisms are unclear. Here, we used a statistical learning paradigm to expose listeners to sequences of speech syllables with fixed transitional probabilities, forming an artificial “language” of trisyllabic words. Following exposure, participants attended to one of two concurrent syllable streams and detected target syllables. Detection performance improved when the attended stream conformed to the statistical structure learned implicitly during exposure, with a larger benefit in the presence of a competing stream than in quiet. In contrast, predictability of the unattended stream had no effect on performance. Electroencephalography revealed that predictable targets elicited earlier parietal P300 “target-recognition” responses and enhanced neural tracking of the attended stream, with additional signatures of predictive processing observed even in the absence of targets. These findings demonstrate that learned statistical regularities enhance listening in noise by enabling predictive, schema-based selection of relevant input. Rather than facilitating automatic segregation of competing sounds, learned lexical schemas support auditory scene analysis through attentional template matching. Our findings establish a direct mechanistic link to the role of prediction in schema-based listening in noise.

**Significance:** Our remarkable ability to isolate a target sound source, such as a person’s voice, in noisy environments is essential for effective communication. This process—termed auditory scene analysis—is known to rely on low-level acoustic regularities, but it is unclear whether learned higher-level regularities, like linguistic structure, also contribute. Combined electroencephalography and behavioral experiments reveal that statistical prediction of upcoming target syllables based on learned syllable-transition probabilities of an artificial language improves attentional selection to a target sound stream in an acoustic mixture. Prediction enhances neural tracking of the attended stream and speeds neural recognition of auditory targets. These findings have implications for auditory training approaches to rehabilitate hearing-impaired individuals who struggle to understand speech in noise.

## 1. Introduction

To listen in noisy environments like crowded streets and cafes, we rely on our brains’ ability to segregate (i.e., separate) a mixture of sound sources into distinct perceptual streams, like speech from background noise. Only when we successfully segregate sources can we direct attention to a target sound source and suppress interfering distractors (Bregman, 1994; Darwin, 1997). Successful attentional selection in turn influences stream formation; thus the processes of segregation and attentive selection interact in a heterarchical manner with each exerting influence on the other (Shinn-Cunningham, 2008). The problem of determining the acoustic content of constituent sources from a sound mixture—also referred to as auditory scene analysis—is mathematically ill-posed. Yet humans are able to successfully navigate it by leveraging statistical regularities in the input sound (McDermott, 2009).

Acoustic regularities like temporal coherence of sound elements across frequency (comodulation) (Elhilali et al., 2009; Hall et al., 1984; Schooneveldt & Moore, 1989; Viswanathan, Bharadwaj, et al., 2021; Viswanathan et al., 2022; Viswanathan, Heinz, et al., 2024) and harmonic structure (McPherson et al., 2022; Rajappa et al., 2023), universally present in natural sounds like speech, provide strong cues for bottom-up (i.e., feedforward or automatic) grouping of different elements of a single sound source into a perceptual object.

Other statistical regularities like linguistic and musical structure must be learned to form a mental “schema” (stored knowledge) of patterns in the acoustic environment (Billig et al., 2013; Bregman, 1994; Denham & Winkler, 2006; Woods & McDermott, 2018). Learned schema cues, like formant transitions in sine-wave speech (Bailey et al., 1977; Remez et al., 1981), melodies (Bey & McAdams, 2002; Dowling, 1973; Woods & McDermott, 2018), and words (Billig et al., 2013), may aid listening in noise.

Although schema-based grouping is thought to rely on predictive modeling of the input sound (Denham & Winkler, 2006; Winkler et al., 2009), a direct mechanistic link between statistical learning, prediction, and auditory scene analysis has not been established. One view is that schemas aid primitive/pre-attentive auditory perceptual organization by supporting sequential grouping of sound elements across time (i.e., streaming) in a bottom-up manner. Thus, when competing sources in a mixture include familiar patterns, source segregation may occur more automatically (Bendixen et al., 2012; Hicks & McDermott, 2024; Johnsrude et al., 2013). Another, competing view is that schemas help listeners focus attention; this idea posits that streams form initially based on acoustic regularities, and that attention selects a target stream by matching sounds in the mixture to the learned patterns of the target (Bendixen et al., 2010; Bregman, 1994).

The term “schema-based segregation” has been used to describe both of these two distinct views, but it fails to distinguish between the essential differences in the two theories. The former account describes a pre-attentive role for schemas in perceptual organization. The second explanation relates to attentional selection using template/pattern matching (e.g., to learned speech-syllable sequences or melodies). The former view predicts that a masker that matches learned schemas will enhance behavioral performance over a more random masker; the latter predicts that only schemas in the attended stream will influence perception. One goal of the current study is to test these competing ideas.

A rich literature on predictive coding has identified electrophysiological correlates of auditory-sequence deviance processing, such as the mismatch negativity response (Casado-Román et al., 2020; Fitzgerald & Todd, 2020; Garrido et al., 2009; Näätänen et al., 1978, 2007; Opitz et al., 2002; Paavilainen et al., 1991; Szalárdy et al., 2020). A parallel language-processing literature has documented electrophysiological correlates of lexical surprisal, e.g., the N400 response (Bakker et al., 2015; Cunillera et al., 2009; Dou et al., 2025; Lau et al., 2009). In addition, studies on statistical learning of sound sequences have quantified neural correlates of such learning using, e.g., event-related potentials (ERPs) like the P300 (the classic target-recognition response (Donchin & Coles, 1988; Picton, 1992; Squires et al., 1973)) and spectral measures of target tracking like intertrial phase-locking value (ITPLV (Tallon-Baudry et al., 1996)) (Abla et al., 2008; Batterink et al., 2015; Batterink & Paller, 2017; Lu et al., 2018; Lu & Vicario, 2014; Pinto et al., 2022). However, neural links between learning, prediction, and auditory scene analysis are under-characterized. A second goal of this study is to explore neural responses to sound streams that either match or do not match learned sequential patterns to gain insight into the neural mechanisms by which learned schemas aid auditory scene analysis.

To answer these questions, we conducted a series of combined behavioral and electroencephalography (EEG) experiments. We exposed participants to sequences of speech syllables with fixed syllable-transition probabilities, following established statistical learning paradigms (McMillan & Saffran, 2016; Saffran et al., 1996, 1997). We hypothesized that as listeners implicitly learned the syllable-transition probabilities of the presented sequence of syllables, they would become better at detecting a target syllable whose occurrence they could predict from the preceding syllables. Post exposure, we tested whether predictability of the attended (experimental paradigm 1) and masker (paradigm 2) syllable streams aids target-syllable detection by independently manipulating the transition probabilities between successive syllables in the attended and masker streams, respectively (see Figure 1).

**Figure 1.**
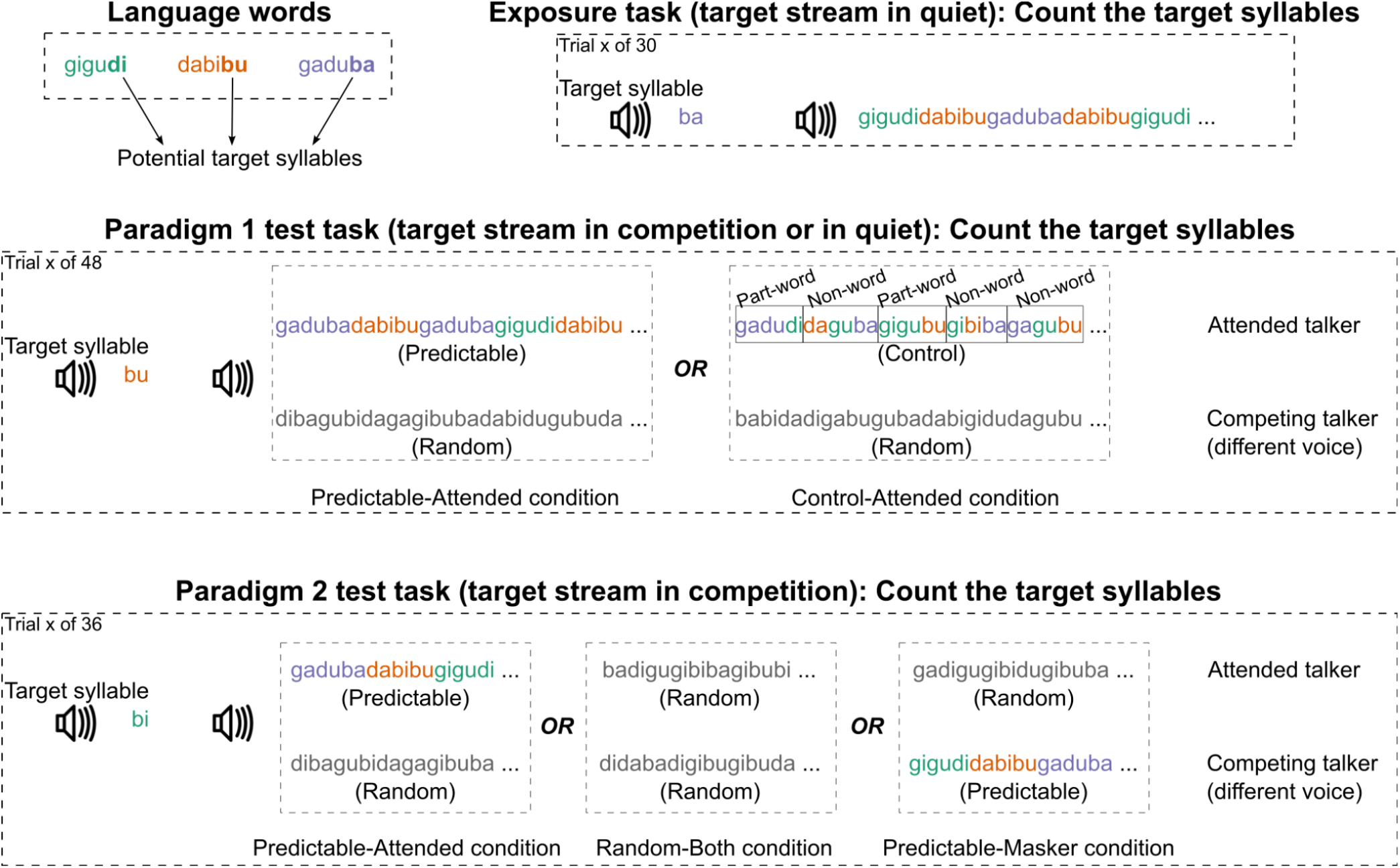
Schematic illustrating the stimuli and experimental tasks. Detailed information is provided in the main Methods text. Briefly, participants underwent ∼20 min exposure to sequences of speech syllables with fixed syllabic transition probabilities (i.e., words from a made-up language). Subjects were tasked with counting the number of times they heard a cued target syllable. Post-exposure, during testing, subjects detected cued target syllables in a target stream either in the presence of a competing syllable stream (spoken by a different talker than the attended talker) or in quiet. By manipulating syllable-transition probabilities in the attended stream (paradigm 1) and the masker stream (paradigm 2) to match those learned implicitly during exposure, we tested how predictability of each stream influences auditory scene analysis.

Our data show that target-syllable detection is enhanced when implicitly learned syllable-transition probabilities in the attended stream predict the target. Importantly, this prediction benefit is greater when the target stream is presented in competing sounds than in quiet, suggesting that syllable-sequence prediction aids schema-based auditory scene analysis. Further, changing the syllable-transition predictability in a simultaneously presented competing syllable stream had no significant effect on performance. These behavioral results support the view that lexical schemas, such as the syllable sequences used in the current study, aid attentional selection of a target speech stream via template matching. EEG data show that the parietal P300 ERP accompanying target-syllable detection occurs earlier when the target is predicted by its preceding syllables. Indeed, the P300 latency is even smaller for predicted targets in a sound mixture than for unpredicted targets in quiet, suggesting that prediction more than overcomes competition from a distracting source. Further, target tracking in the brain, as indexed by ITPLV, is stronger in the predictable condition than in the unpredictable condition. Finally, we also observe purely endogenous (in the absence of target detection) neural correlates of predictive processing. Taken together, our findings support the interpretation that statistical prediction shapes schema-based attentional selection of a target sound stream; prediction enhances target tracking in the brain and leads to earlier neural recognition of auditory targets.

## 2. Results

### 2.1. Sequence prediction aids schema-based auditory scene analysis

Subjects were exposed to sequences of speech syllables with consistent syllable-transition probabilities. Specifically, sequences of three three-syllable-long made-up words were presented, following established statistical learning paradigms (McMillan & Saffran, 2016; Saffran et al., 1996, 1997). Afterwards, subjects performed a target-syllable detection task where they listened for a particular syllable within a target stream. We contrasted presenting the target stream alone versus in the presence of a similar, competing stream. At the start of each individual trial, a randomly selected syllable was played to cue the subjects as to the target syllable for that trial. Subjects were tasked with detecting and counting the number of times they heard that target syllable in the attended stream while ignoring any instances of that syllable in a competing syllable stream spoken by a different talker voice (paradigm 1; see Figure 1).

The target syllable was always the third syllable of a triplet (sequence of three syllables) within the target stream. The attended stream was either predictable, consisting of words from the artificial language that subjects heard during exposure (Predictable-Attended condition), or a control stream, where syllable transitions did not match those presented in the exposure period and the target syllable thus could not be predicted by the preceding syllables (Control-Attended condition). The control stream was made up of syllable triplets that were either “part-words” (where the first two syllables of a triplet were from a word in the artificial language, but the third syllable did not match the expected final syllable, but was chosen as one of the other possible final-position syllables from other words in the language) or “non-words”. In “non-words,” syllable positions were constrained such that the ith syllable of each triplet was always one of the syllables appearing in the ith syllable of a triplet in the language words, but neither the first-to-second nor second-to-third syllable transition matched that in the language. The ignored stream was always a completely random sequence of the same syllables used in the artificial language.

In-lab data (experiment 1 of paradigm 1) show that overall, listeners performed more accurately when targets were predictable than in the control condition, both in the presence of a competing stream and in quiet (Figure 2A). The prediction benefits (percent changes in performance between the Control-Attended and Predictable-Attended conditions) on percent trials correct in competition and in quiet were consistently positive (Figure 2B). Consistent with this, the Size of Error (SOE; the square root of the mean squared difference between reported and actual target-syllable count across incorrect trials) was smaller in the Predictable-Attended condition than the Control-Attended condition (Figures 2C and 2D).

**Figure 2.**
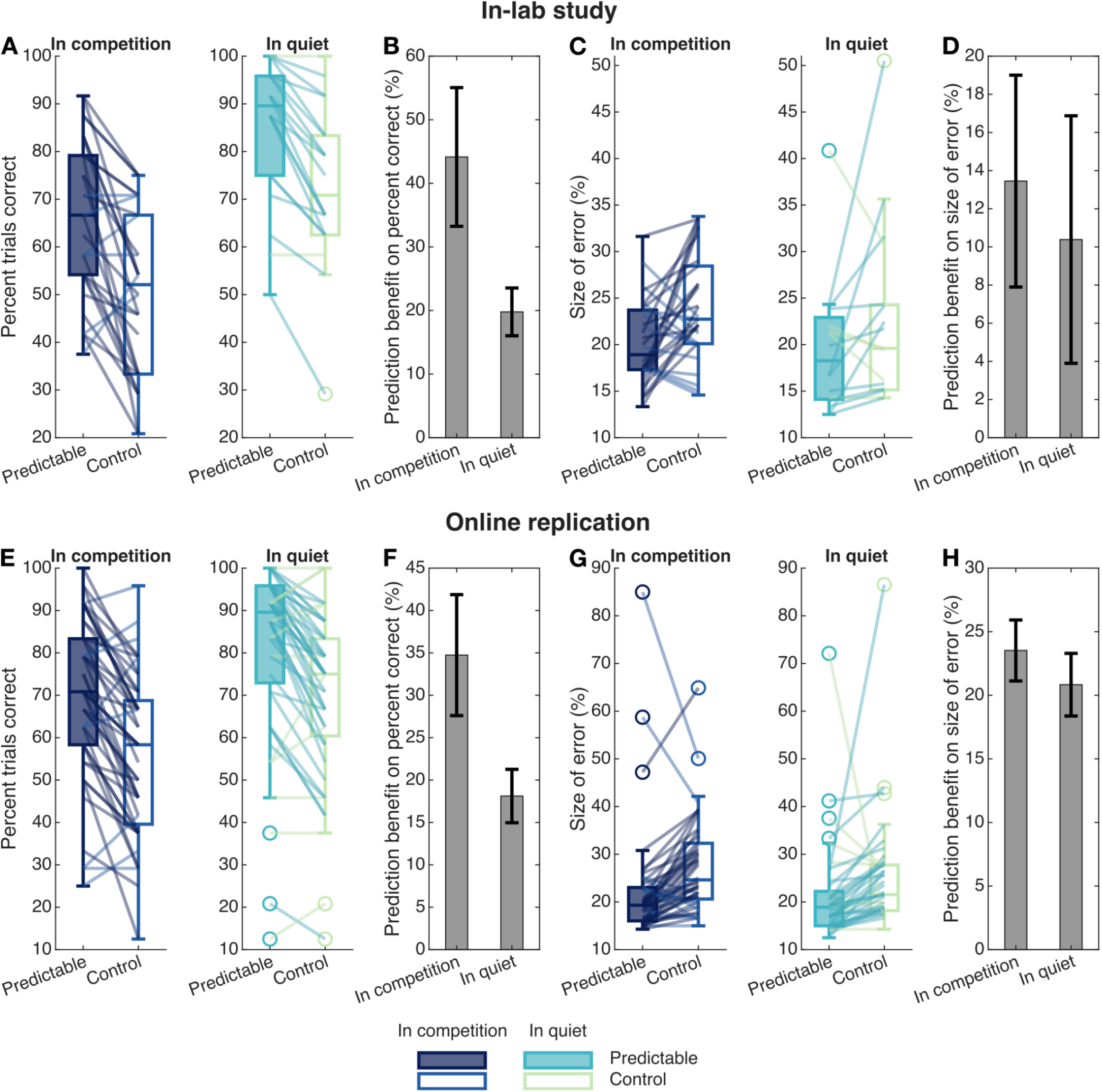
Behavioral effect of attended-stream predictability for target stream in competition and in quiet: paradigm 1 test-stage data for experiment 1 (in-lab study; panels A through D) and experiment 2 (online replication study; panels E through H). (A, E) Percent trials correct (percent trials in which the number of target syllables reported match the number presented), (B, F) prediction benefit on percent trials correct (within-subject percent change in performance between the Control-Attended and Predictable-Attended conditions), (C, G) size of error (SOE; square root of the mean squared error between the number of target syllables reported and presented across incorrect trials, expressed as a percentage relative to the average number of targets presented across incorrect trials), and (D, H) prediction benefit on SOE. In panels A, C, E, G, boxplots show the median (central line), the 25th and 75th percentiles (bottom and top edges of the box), the most extreme data points not considered outliers (whiskers), and the outliers (‘o’ symbol) across subjects; lines connect individual subject percentages for Predictable-Attended and Control-Attended conditions, with the line color reflecting whether performance was better in the predictable condition or the control condition. In panels B, D, F, H, bar plots show the mean and standard error.

Statistical tests confirm these observations. The prediction benefit on percent trials correct is significantly greater than zero for both a target stream played with a competing syllable stream and in quiet (within-subject permutation testing; p < 1e-5 for both masking conditions). The prediction benefit on SOE is significantly greater than zero in competition (p = 0.017) but not in quiet (p = 0.093). These results show that target-syllable detection is easier when learned knowledge of between-syllable transition probabilities in the attended stream predicts an upcoming target. Thus, contextual prediction based on learned schemas aids syllable perception both in competition and in quiet.

A permutation test directly comparing the prediction benefit on percent trials correct across masking conditions (Figure 2B) further showed that the prediction benefit on percent-correct performance was significantly larger in competition than in quiet (between-subject permutation testing; p = 0.028), confirming an interaction between predictability and listening in noise. Taken together, these results suggest that predictive processing of learned schemas aids auditory scene analysis on top of providing a benefit for detecting upcoming syllables within a stream.

An online experiment (experiment 2 of paradigm 1) replicated these in-lab behavioral findings. The online experiment used a fully within-subjects design, with each subject performing both masking conditions (in competition, and in quiet) in addition to both contextual conditions (Predictable-Attended, and Control-Attended). As in the in-person study, the mean prediction benefit was positive for both target streams presented in competition and in quiet (Figure 2E,F,G,H), and the mean benefit was larger in competition than in quiet (Figure 2E,F).

Statistical tests supported these observations. The prediction benefit on percent trials correct is significantly greater than zero both in competition and in quiet (within-subject permutation testing; p < 1e-5 for both masking conditions). Furthermore, the prediction benefit on percent trials correct is greater in competition compared to in quiet (within-subject permutation testing; p = 0.0056). The prediction benefit on SOE is also significantly greater than zero for both the in-competition (p = 5e-5) and in-quiet (p = 4e-5) conditions; however, the prediction benefits on SOE do not significantly differ across these two masking conditions (p = 0.25). These online data support our hypothesis that statistical prediction based on learned schemas improves perceptual separation of a target from a masker and/or attentional focus to the target stream, thereby aiding syllable perception in the presence of competing sounds.

### 2.2. Neural recognition occurs earlier for predictable compared to unpredictable target syllables

Target syllables within the attended stream reliably evoke a P300 ERP (a positive deflection roughly 300 ms after the target syllable that is typically observed when an unlikely target is detected (Picton, 1992; Polich, 2007)) with a parietal scalp distribution, consistent with recognition of this behaviorally important syllable; no such response is present when the target is absent (Figure 3A,B) (Halgren et al., 1998; Ji et al., 1999; Vaughan & Ritter, 1970; Wronka et al., 2012). Moreover, no P300 response is observed for target syllables in the ignored/masker stream, consistent with listeners successfully ignoring the masker while attending to the cued target stream.

**Figure 3.**
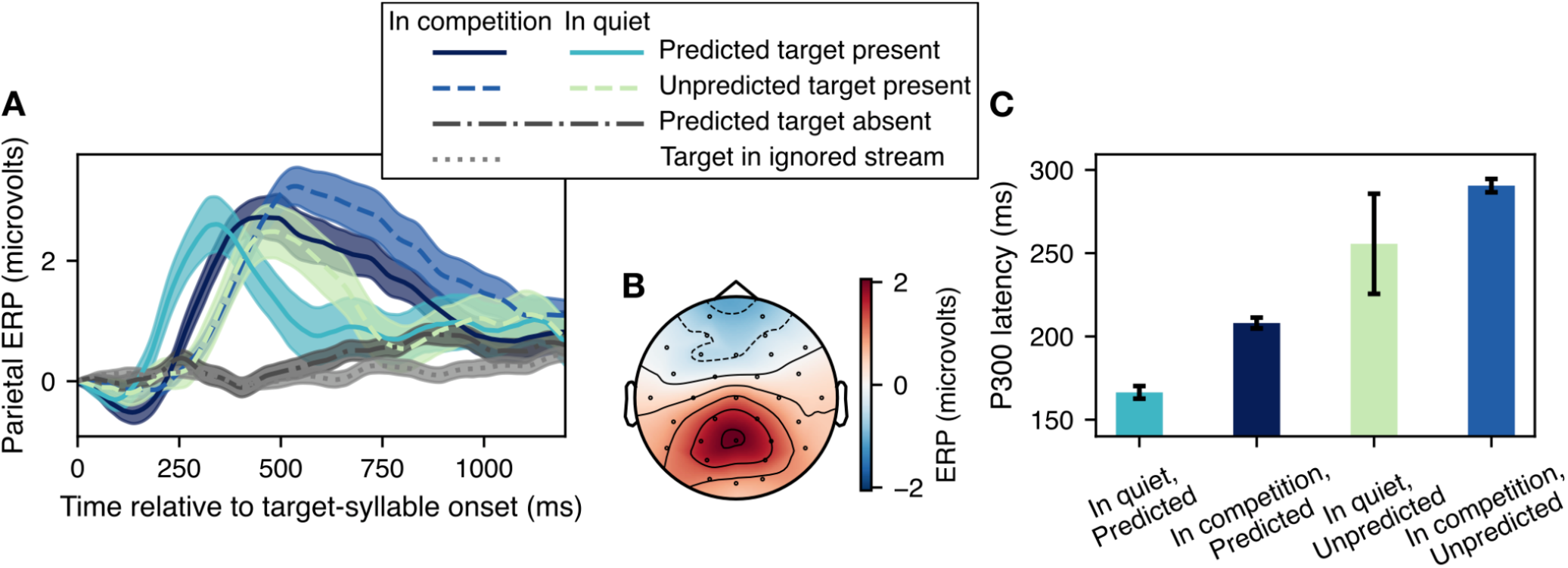
P300 ERPs (paradigm 1 test-stage EEG data; mean and standard error across subjects). (A) Average ERP time course in parietal electrodes (Pz, CP1, CP2, P3, and P4) for third-syllable onsets in the attended stream in the different contextual (Predictable-Attended or Control-Attended/Unpredictable) and masking (target stream in competition or in quiet) conditions, and for the cases when the third syllable is the target and when it is not the target. Data for the case when the target syllable is predicted but absent are pooled over both masking conditions. Data are also shown for the case when the target syllable occurs in the ignored stream. (B) Scalp topomap of the P300 target-recognition response averaged across conditions, subjects, and time (between 0 and 300 ms post the P300 latency); the topomap illustrates the parietal scalp distribution of the P300. (C) Bar plots showing the earliest-response latency of the parietal P300 ERP in the different contextual and masking conditions, sorted by mean latency.

The latency of the P300 target-recognition response varies with predictability of the target and whether or not there is a competing masker stream (Figure 3C). The P300 (specifically, the earliest response; see Materials and methods: P300 latency computation) occurs earlier when the preceding syllables predict the target syllable (Predictable-Attended condition) compared to when they do not (Control-Attended condition) [t(28) = -88.44, p < 2.2e-16 for target stream in competition; t(19) = -13.52, p = 1.7e-11 for target stream in quiet]. Induced alpha and beta power show an analogous difference in timing (see Supplementary Figure S1).

Importantly, the P300 latency for predicted target syllables in competition is even smaller than that for unpredicted target syllables in quiet [t(19.3) = -6.87, p = 6.85e-7]. Because background noise can delay target processing (Ratcliff & McKoon, 2008), this result suggests that predictability of sound sequences affords a processing speed benefit for syllables presented with competing sounds. Although we observe a small target-syllable-evoked N100 ERP (a negative deflection roughly 100 ms after the evoking syllable) in the Predictable-Attended condition, we do not observe a clear N100 response in the Control-Attended condition (Supplementary Figure S2).

### 2.3. Lexical schemas support attentional selection of the target stream (via template matching to learned words) rather than aiding pre-attentive auditory scene analysis

Prior studies raise the possibility that masker schemas like sound textures (Hicks & McDermott, 2024) and a familiar voice (Johnsrude et al., 2013) may aid perceptual organization. To test whether this may also be the case for learned syllable sequences, we conducted a follow-up fully online experiment (paradigm 2; see Figure 1). The procedure in the exposure stage of paradigm 2 was identical to that of paradigm 1. However, in the test stage of paradigm 2, we manipulated the predictability of the masker syllable stream independently of that of the attended syllable stream across different trials. If learned schemas aid pre-attentive scene analysis, performance should be better when the distracting stream is made up of triplets from our learned language compared to when the syllable triplets are random. Conversely, if the statistical structure of the distractor has no effect on performance, it suggests that schemas serve primarily to aid attentional focus, not to bolster automatic auditory scene analysis.

In paradigm 2, both target and competing/masker streams were made up of sequences of the same syllables; however, which of the streams followed the implicitly learned structure of the artificial language varied across conditions. The paradigm used a fully within-subjects design where each subject performed the following experimental conditions: (i) Predictable-Masker, in which the masker stream consisted of words from the artificial language while the attended stream consisted of a random, unstructured syllable sequence, (ii) Random-Both, in which the attended and masker streams each consisted of an independently drawn random syllable sequence, and (iii) Predictable-Attended, in which the attended stream consisted of words while the masker stream consisted of a random syllable sequence.

Performance was consistently better in the Predictable-Attended condition than in the two conditions with random target streams (Predictable-Masker and Random-Both conditions) (see Figure 4), replicating our finding that attended-stream syllable-sequence predictability aids target detection. Moreover, performance in the Predictable-Masker and Random-Both conditions was nearly identical, both when bottom-up segregation based on acoustic cues was relatively easy (the target and masker streams were spoken by talkers with different genders; experiment 1 of paradigm 2; Figure 4A,B,C,D) and when it was difficult (the two streams were spoken by female talkers differing only in voice pitch and by 50 Hz; experiment 2 of paradigm 2; Figure 4E,F,G,H).

**Figure 4.**
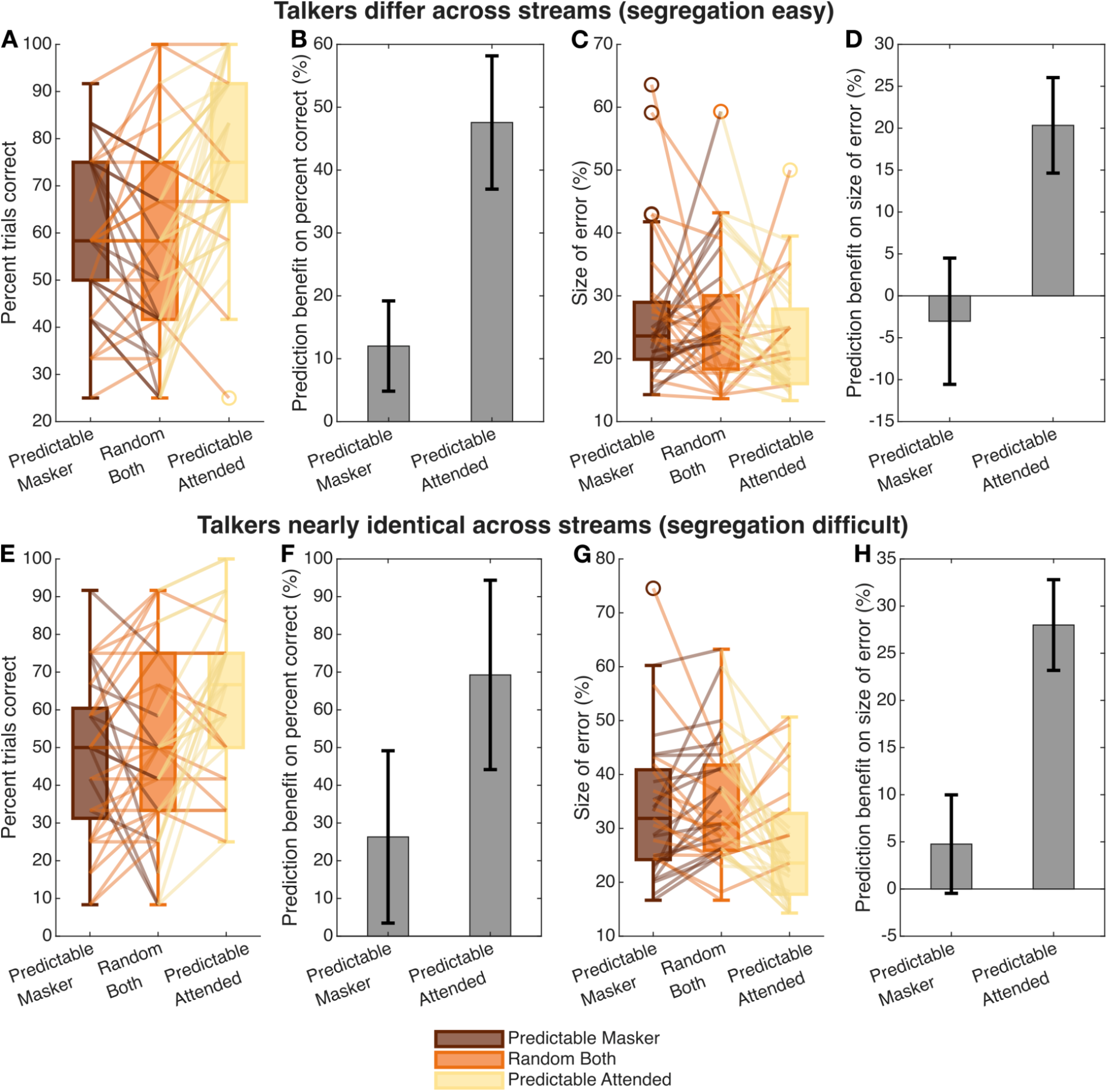
Behavioral effects of masker- and attended-stream predictability: paradigm 2 test-stage data for experiment 1 (segregation easy; panels A through D) and experiment 2 (segregation difficult; panels E through H). (A, E) Percent trials correct (percent trials in which the number of target syllables reported match the number presented), (B, F) prediction benefit on percent trials correct for when the masker and attended stream are each predictable (within-subject percent changes in performance between the Random-Both and Predictable-Masker conditions, and between the Random-Both and Predictable-Attended conditions, respectively), (C, G) size of error (SOE; square root of the mean squared error between the number of target syllables reported and presented across incorrect trials, expressed as a percentage relative to the average number of targets presented across incorrect trials), and (D, H) prediction benefit on SOE for the cases when the masker and attended streams are each predictable. In panels A, C, E, G, boxplots show the median (central line), the 25th and 75th percentiles (bottom and top edges of the box), the most extreme data points not considered outliers (whiskers), and the outliers (‘o’ symbol) across subjects; lines connect individual subject percentages for Predictable-Masker, Random-Both, and Predictable-Attended conditions, with the line color reflecting the condition in which performance was better. In panels B, D, F, H, bar plots show the mean and standard error.

Confirming these observations, the prediction benefit was significantly greater than zero when the attended stream was predictable, for both the easy segregation condition (within-subject permutation testing; p = 3e-5 for percent trials correct and p = 0.0078 for SOE) and the difficult segregation condition (p = 7.4e-4 for percent trials correct and p = 1.3e-4 for SOE). However, there was not a statistically significant benefit of masker predictability for either the easy segregation condition (p = 0.15 for percent trials correct and p = 0.64 for SOE) or the difficult segregation condition (p = 0.55 for percent trials correct and p = 0.23 for SOE). This is especially notable given that there was more room for masker schemas to improve auditory scene analysis in the difficult segregation condition, where performance was farther from ceiling.

Behavioral error patterns in target-syllable detection for target syllables in competition are also revealing. On average, across all conditions listeners undercounted target syllables (Figure 5). However, the magnitude of this undercounting was significantly smaller when the target stream was predictable compared to when it was unpredictable, regardless of the masker-stream predictability (within-subject permutation testing; for Control-Attended vs. Predictable-Atttended in paradigm 1, p < 1e-5 for both experiments 1 and 2; for Predictable-Masker vs. Predictable-Atttended in paradigm 2, p = 5.5e-4 for experiment 1 and p = 0.021 for experiment 2; for Random-Both vs. Predictable-Atttended in paradigm 2, p = 6.8e-4 for experiment 1 and p = 0.0047 for experiment 2).

**Figure 5.**
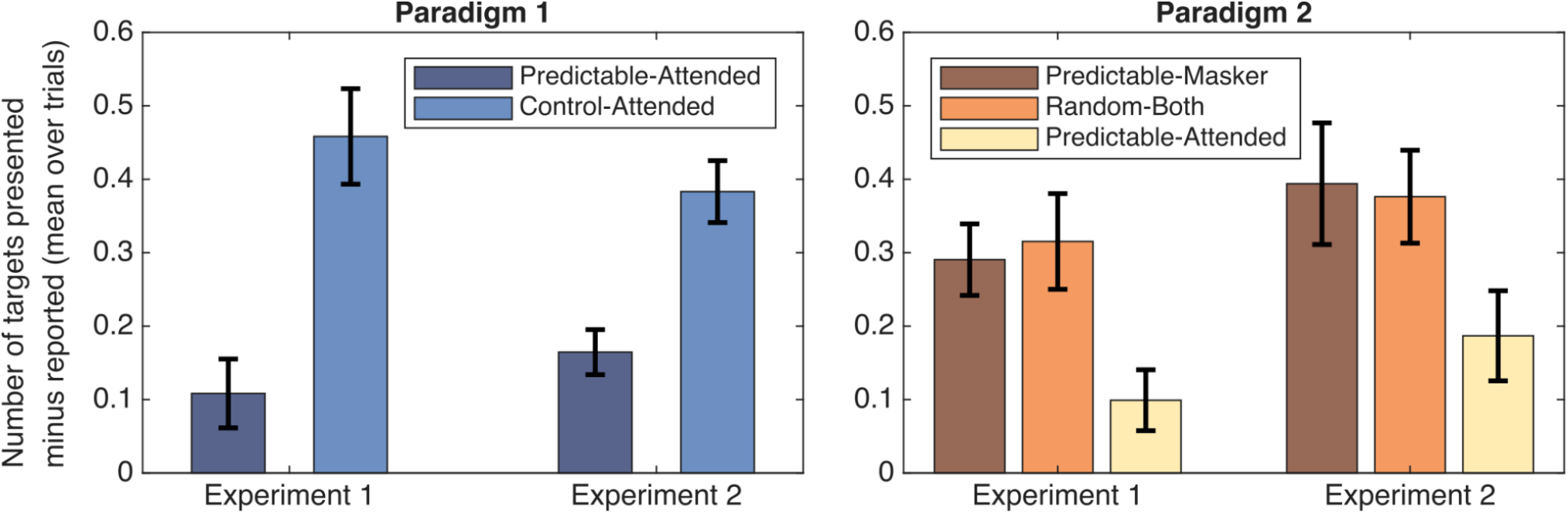
Behavioral error patterns for target stream in competition (test-stage data; mean and standard error across subjects). Bar plots show the differences between the number of targets presented and reported, averaged over trials, for the different contextual conditions in each paradigm and experiment.

Thus, errors arose because listeners occasionally missed target syllables in the attended stream. Our results show that syllable-sequence predictability improves performance—by reducing the number of missed target syllables within the target stream. This finding supports the interpretation that prediction helps with maintaining sustained attention on the target stream and that predictive processing may help avoid missed target detection by cueing the attentional system to the impending occurrence of a target.

### 2.4. Predictability strengthens neural tracking of the target stream

Evoked neural responses are stronger, as measured by the intertrial phase-locking value (ITPLV), in the Predictable-Attended condition compared to the Control-Attended condition (Figure 6). Specifically, the ITPLV at the first harmonic of the triplet rate (the triplet rate is ∼0.72 Hz, and its first harmonic is ∼1.45 Hz) and the syllabic rate (2.17 Hz) are both significantly larger in the Predictable-Attended condition than the Control-Attended condition for target stream in competition [t(28) = 3.26, p = 0.0015 and t(28) = 3.9, p = 0.00028, respectively]. This difference in neural response strength occurs despite the fact that the stimulus power does not differ between these contextual conditions at either of these frequencies (Supplementary Figure S3 shows the modulation spectra of the stimuli in paradigm 1 for the different contextual conditions). This result suggests that prediction enhances neural tracking of the target stream. Moreover, we observe enhanced neural tracking in the predictable condition at timescales longer than the duration of individual syllables (i.e., at ∼1.45 Hz), in line with the multi-syllabic structure of the imposed predictability.

**Figure 6.**
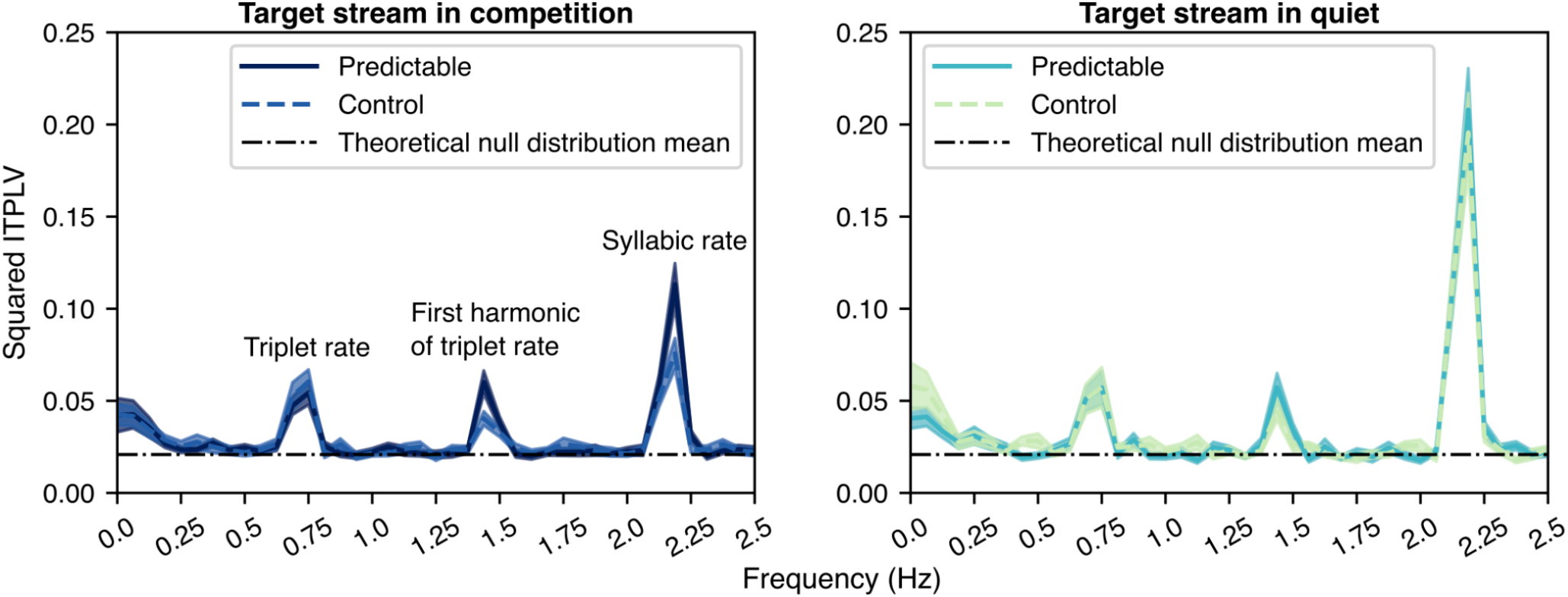
Squared intertrial phase-locking value (ITPLV) spectra for target stream in competition (left plot) and in quiet (right plot) (test-stage EEG data from paradigm 1; mean and standard error across subjects). The ITPLV spectra are averaged over all 32 EEG channels, and shown separately for the Predictable-Attended and Control-Attended conditions. The theoretical null distribution mean for the squared ITPLV measure is also shown.

Attention to a stream is known to cause that stream to evoke larger neural responses than when that same acoustic input is being ignored (Choi et al., 2014; Donchin & Cohen, 1967; Hillyard et al., 1973; Sheatz & Chapman, 1969; Viswanathan et al., 2019). Thus, these results are consistent with the interpretation that predictive processing of learned schemas improves the efficacy of selective attention. Supporting this explanation, although we see a large contextual effect on ITPLV in competition, we see only a small, albeit statistically significant, effect in quiet at the first harmonic of the triplet rate [t(19) = 2.054, p = 0.027] and no significant effect in quiet at the syllabic rate [t(19) = 1.29, p = 0.11] (Batterink & Paller, 2017; Pinto et al., 2022).

### 2.5. Neural correlates of predictive processing appear even when the target is absent

The Control condition included part-words, where the first two syllables of a triplet match the first two syllables of a word in our learned language (but the final syllable differs from expectations), as well as non-words, where none of the syllable transitions obey the learned language. In both masking conditions, ERPs to part-words differed significantly from non-words starting with the same initial syllable (Figure 7A). Specifically, ERPs to target-predicting part-words and non-words differ significantly in the time period between the onsets of the second and third syllables [t(28) = 2.69, p = 0.012 in competition; t(19) = 2, p = 0.0602 in quiet]. Because this effect is not directly related to the detection of the cued target, it can be unambiguously attributed to an endogenous influence of schema-based predictive processing—a clear neural correlate of predictive processing based on our learned language. The scalp topomap of this ERP difference (Figure 7B) does not match the topomaps of either the P300 (Figure 3B), the well-established N100 (Viswanathan, Bharadwaj, et al., 2021), or the mismatch negativity response (Giard et al., 1990), which suggests that the neural generators underlying the endogenous predictive-processing effect seen in Figure 7 may be distinct from those underlying these classic ERPs (see Discussion).

**Figure 7.**
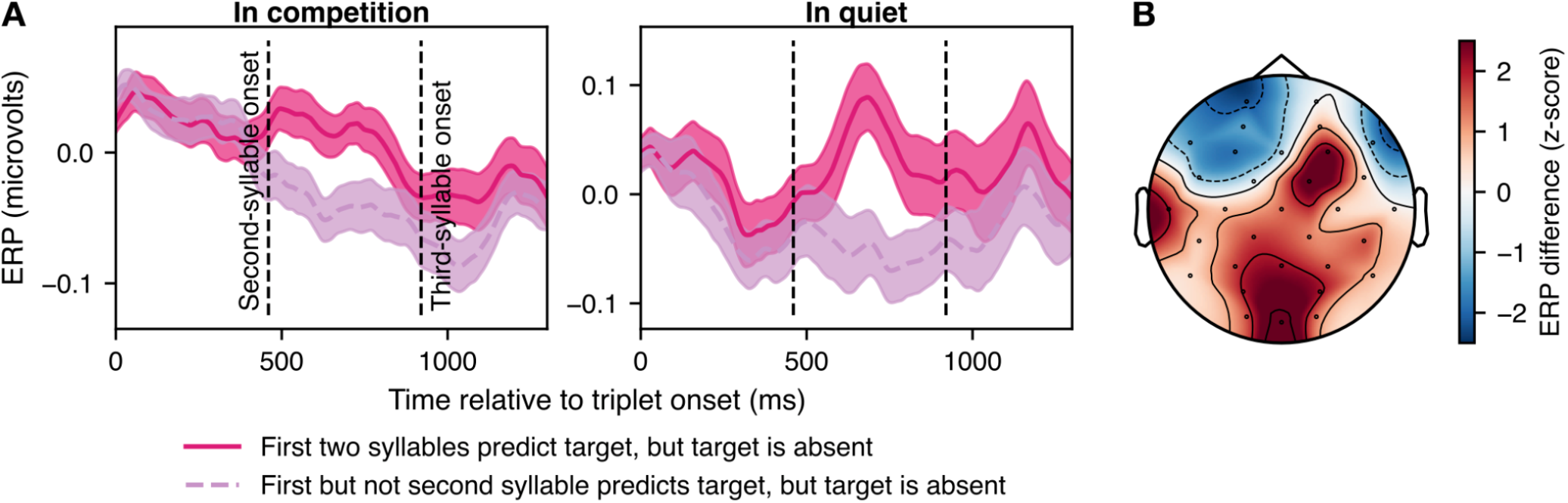
Comparison of ERPs for part-words that predict an upcoming target (the first two syllables match the syllables in the word that contains a target third syllable) and non-words where the first (but not the second) syllable predicts the target (paradigm 1 test-stage EEG data from Control-Attended trials in which the target syllable is not presented). (A) ERP timecourses locked to the onset of triplets (sequences of three syllables) for target stream in competition and in quiet (mean and standard error across subjects). Time zero corresponds to the onset of the first syllable; the onsets of the second and third syllables are marked. Note that the ERPs shown for each masking condition are channel-weighted by the corresponding average (across subjects and time between the onsets of the second and third syllables) scalp topomaps of the ERP differences between target-predicting part-words and non-words. (B) Average (across subjects and time between the onsets of the second and third syllables) scalp topomap of the ERP difference between target-predicting part-words and non-words (data pooled over conditions with and without a competing stream).

## 3. Discussion

Our behavioral and EEG data show that sequence prediction based on learned lexical schemas aids auditory scene analysis, specifically, attentional selection of a target sound stream in an acoustic mixture. Predictability strengthens neural tracking of the target stream and shortens the latency of neural recognition of auditory targets, both of which help listening in noisy settings. Our findings provide a direct, mechanistic link between statistical learning, predictive processing, schema-based auditory scene analysis, and selective attention.

Although some prior studies have suggested that lexical knowledge may support build-up of streaming (sequential grouping) and detection of a subsequent target sound, these studies—unlike the current study—did not examine whether there is an interaction between lexical predictability and the degree to which a competing stream interferes with processing of a target stream (Billig et al., 2013; Szalárdy et al., 2020). Further, past data do not disambiguate between the pre-attentive and attentive accounts of schema-based auditory scene analysis.

More generally, the term “schema-based segregation” has been used in the literature to refer to both the perceptual organization and attentional selection accounts of schema-based auditory scene analysis. However, varying predictability in the masker sequence of syllables had no effect on performance, even in a relatively hard segregation condition (Figure 4). This finding contradicts the pre-attentive perceptual organization view of schema-based auditory scene analysis: if familiarity of syllable sequences aided pre-attentive perceptual organization, performance would be better when the masker stream was structured. Instead, our results suggest that streams following learned lexical schemas enhance attentional selection of the target stream, not primitive perceptual organization. This finding is consistent with prior work using interleaved melodies overlapping in pitch range that did not find any effect of knowledge of the background melody on recognition of a target melody (Dowling, 1973). It is also in line with prior EEG studies that used neural tracking measures to suggest that a sequence of syllables is not grouped into multisyllabic words when attention is not directed towards that stream (Ding et al., 2018). Some studies have demonstrated that predictability of competing source elements like tones, sound textures, and familiar voices do improve perception of a simultaneously presented target source (Bendixen et al., 2012; Hicks & McDermott, 2024; Johnsrude et al., 2013). However, these experiments did not manipulate the predictability of the sequence order of categorizable, countable elements making up the competing source, but rather the statistics of its overall spectrotemporal content. Thus, it is possible that different mechanisms may be involved in schema-based auditory scene analysis when the learned structure depends on analyzing the sequential order of distracting-source events versus the overall sound quality of the competing source. While the latter class of schemas may support primitive perceptual organization, the former may only play a role in selective attention.

Our finding that the P300 target-recognition response occurs earlier for predicted target syllables (Figure 3) is in line with prior results that the P3b ERP latency is shorter for target tones that are predicted by preceding tones, matching a learned tonal pattern (Fogelson et al., 2009). It is also consistent with the shorter behavioral reaction times documented for predicted compared to unpredicted targets (Batterink et al., 2015; Fogelson et al., 2009). However, our results extend this previous work by demonstrating that the P300 latency for predicted targets *in competition* is even shorter than that for unpredicted targets *in quiet*. This result suggests that predictability of sound sequences speeds target recognition enough to more than offset slowed processing in the presence of background noise (Ratcliff & McKoon, 2008). Indeed, drift diffusion models suggest that evidence accumulation is slower when task difficulty is greater, delaying decision making. Our results provide empirical evidence that predictive processing based on prior knowledge speeds processing and can offset such delays (Knill & Pouget, 2004). Note that although we observe an effect of contextual condition on the P300 latency, we do not observe any such effects on the P300 amplitude (Batterink et al., 2015).

By parametrically varying target predictability (using language words, part-words, and non-words), we quantified neural correlates of purely endogenous predictive processing, i.e., processing of mismatch between pre-learned stimulus statistics and the presented stimuli when a predicted target was actually absent. Future studies should use high-density EEG or magnetoencephalography (MEG) and source localization techniques to explore the neural bases of the complex scalp topography pattern corresponding to this endogenous predictive-processing effect (Figure 7).

We had subjects perform a target-syllable counting task during exposure rather than having them passively listen to stimuli because we wanted to promote engagement. However, in line with established statistical learning paradigms that we adopted here (McMillan & Saffran, 2016; Saffran et al., 1996, 1997), participants were not told about the syllable-transition probabilities present in the stimuli. Because of this, and because knowledge of stimulus statistics was not necessary to do the target-syllable counting task, learning during exposure was implicit (Reber, 1967). Our data show modest performance improvements after the first few trials of exposure, demonstrating that implicit syllable-sequence learning occurs rapidly (Supplementary Figure S4), as documented in previous studies (Batterink & Paller, 2017; Saffran et al., 1999). Crucially, we also show that implicit syllable-sequence learning transfers to perceptual outcomes in background noise (Figures 2, 3); this novel finding has implications for aural training. Future work should explore whether the behavioral and neural prediction benefits for cocktail-party listening documented in this study (Figures 2, 3, 5, 6) differ when learning during exposure is explicit rather than implicit. Further, because statistical learning may occur during passive listening, i.e., when subjects are not performing a task (Batterink & Paller, 2019; Saffran et al., 1997), and also when there is acoustic competition (McDermott et al., 2011; McMillan & Saffran, 2016), future work should examine how these different learning conditions influence schema-based auditory scene analysis. Such investigations have the potential to inform clinical practice, e.g., for rehabilitation of hearing-impaired patients (Dornhoffer et al., 2024; Sweetow & Palmer, 2005).

## 4. Materials and Methods

### 4.1. Overview of approach

Participants first underwent exposure to sequences of speech syllables with consistent syllabic transition probabilities (i.e., words from a made-up language). Post-exposure, subjects detected cued target syllables either in the presence of a competing syllable stream or in quiet. We tested how predictability of the attended and masker streams each influences auditory scene analysis by (1) manipulating the syllable-transition probabilities in the attended stream to match those learned implicitly during exposure (paradigm 1), and (2) independently manipulating the syllable-transition probabilities in both streams (paradigm 2). Paradigm 1 included one in-lab experiment and one online replication experiment, whereas paradigm 2 included two fully online experiments.

### 4.2. Participants

In-lab EEG and behavioral data were collected from 58 human subjects (36 female) over the course of one visit to the lab. Subjects were recruited from the Pittsburgh community. All subjects were aged 18–60 years, were native speakers of North American English, and had normal hearing (audiometric thresholds of 25 dB HL or better up to 8 kHz in our hearing screening), no persistent tinnitus (by self report), and no known neurological disorders (by self report). Subjects participated for course credit or pay. All study procedures were approved by the Carnegie Mellon University institutional review board (IRB). Subjects provided informed consent in accordance with these procedures.

Online behavioral data were collected on a web-based psychoacoustics platform (Mok et al., 2024) from anonymous subjects recruited using Prolific.co. 49 subjects performed the online replication experiment (experiment 2) of paradigm 1, 40 subjects performed the first experiment of paradigm 2, and 40 performed the second experiment of paradigm 2. The subject pool was restricted using a screening method developed by Mok et al. (Mok et al., 2024). The screening method contained three parts: (i) a core survey that was used to restrict subjects based on age to 18–55 years (to exclude significant age-related hearing loss); to ensure they were US/Canada residents, US/Canada born, and native speakers of North American English (because North American speech stimuli were used); and to ensure they had no history of hearing and neurological diagnoses or persistent tinnitus; (ii) headphone/earphone checks; and (iii) a speech-in-babble-based hearing screening. Subjects who passed the screening were invited to participate in the main study, and when they returned, headphone/earphone checks were performed again. All subjects had completed at least 40 previous studies on Prolific and had >90% of them approved. These procedures were validated in previous work, where they were shown to successfully select participants with near-normal hearing status, attentive engagement, and stereo headphone use (Mok et al., 2024). Subjects provided informed consent in accordance with remote testing protocols approved by the University of Pittsburgh IRB.

### 4.3. Stimuli and experimental design

#### 4.3.1. Stimuli

We created 12 artificial languages in total and used a different/non-overlapping set of four artificial languages each in paradigm 1 (the in-lab and replication experiments of this paradigm used the same four languages), the first experiment of paradigm 2, and the second experiment of paradigm 2. Each artificial language was composed of three three-syllable made-up words (hereafter referred to as “language words”). Within each artificial language, the three three-syllable words in the language dictionary were created using the speech syllables, /ba/, /da/, /ga/, /bi/, /di/, /gi/, /bu/, /du/, /gu/, chosen randomly without replacement (see Figure 1).

Paradigm 1 used two sets of syllables, one from a female and one from a male native speaker of North American English. Paradigm 2 used the same syllable tokens as paradigm 1, and, in addition, also used tokens from the female talker with the voice pitch shifted downwards by 50 Hz. Hereafter, we will refer to the original and the pitch-transposed female talker voices as the first and second female talkers, respectively.

Syllables were 400 ms long, and ramped on with a rise time of 0.01 s and ramped off with a fall time of 0.01 s. Syllables were concatenated with a 60-ms silence between successive syllables to create language words and sequences of words (or of syllables, more generally). This yielded a syllabic rate of ∼2.17 Hz and a triplet (sequence of three syllables) frequency of ∼0.72 Hz. All tasks in the exposure and testing phases of both in-person and online experiments presented stimuli diotically.

#### 4.3.2. Exposure

In both paradigms 1 and 2, each subject first underwent ∼20 min of exposure across 30 trials to a syllable stream consisting of words from one of the artificial languages, as in established statistical learning paradigms (McMillan & Saffran, 2016; Saffran et al., 1996, 1997). Each trial lasted ∼27 s and presented 20 language words in total; within each trial, consecutive words did not repeat. Note that each subject was exposed to only one artificial language and an approximately equal number of subjects performed each artificial language.

To promote engagement, subjects were asked to silently count the number of times they heard a particular target syllable in each trial. Subjects were presented with a visual cue that displayed the target syllable and either the gender (in paradigm 1 and the first experiment of paradigm 2) or pitch (low versus high; in the second experiment of paradigm 2) of the target voice on the computer screen, as well as an auditory cue in which the target syllable was spoken in the voice of the speaker to be attended. The speech stimuli started ∼4 seconds after these cues were presented. Within a given trial, the cued target syllable was presented at most 8 times. At the end of each trial, subjects used a graphical user interface to report their target-syllable count (0, 1, …, and 8). Following their response, an onscreen message revealed the correct number of target syllables in the attended stream.

Target syllables were always the last (third) syllable of a language word; thus, each artificial language had only three possible target syllables. The three candidates for target syllable were presented an equal number of times and interleaved across trials. We predicted that as subjects learned the syllable-transition probabilities of the presented syllables, they would become better at detecting a particular target syllable because they would be able to predict the target syllable from the sequence of preceding syllables (e.g., predict that /di/ always follows /gi/-/gu/ and /bu/ always follows /da/-/bi/ for the artificial language shown in Figure 1). Note that subjects were never told that there was any learning or prediction involved in the study, or that the stimuli used words from an artificially constructed language; thus, the learning underwent by listeners during the exposure stage was implicit.

In paradigm 1, we exposed half of the subjects to the first female talker, and the other half to the male talker. In paradigm 2, in the first experiment, all subjects were exposed to both the first female talker and the male talker alternated across trials; in the second experiment of paradigm 2, all subjects were exposed to both female talkers alternated across trials.

Post-exposure, subjects performed testing, which differed between paradigms 1 and 2.

#### 4.3.3. Paradigm 1 testing

In paradigm 1, 37 in-lab subjects performed a target-stream-in-competition task and 21 in-lab subjects performed a target-stream-in-quiet task. Furthermore, 49 online subjects performed both tasks, which allowed for a fully within-subject quantification of effects.

To achieve this within-subjects design, we posted two separate studies on Prolific.com for each artificial language. One study contained an exposure task followed by a task where the target stream was presented with a simultaneous, competing stream spoken by the other talker. The other study contained the same exposure task followed by a task where the target stream was presented in quiet. Subjects who completed one of the two online studies were invited to participate in the other study; the order of the two studies was counterbalanced across subjects. Note that in this design, each participant in the online experiment who opted to perform both studies performed the exposure task twice. We only included data from the 49 participants who chose to perform both studies.

In each trial of the target-stream-in-competition task, subjects were tasked with detecting and silently counting the number of times they heard a particular target syllable (embedded in a sequence of syllables), cued as in the exposure. At the same time, a competing syllable sequence was played (an unstructured/random sequence of the same nine syllables) at the same root-mean-square sound level (0 dB signal-to-noise ratio). Just as in the exposure training, subjects reported the count of target syllables in the target stream at the end of the trial, following which an onscreen message revealed the correct number of target syllables in the attended stream. As before, target syllables were always the last syllable of a triplet, and the three candidates for target syllable occurred an equal number of times in each condition and were randomly interleaved across trials.

The attended and masker streams were spoken by different talker genders. The same two voices that were used during the exposure stage of paradigm 1, i.e., the first female talker and the male talker, were also used during testing. Both talkers were used for the attended stream, with the attended-talker gender randomly interleaved across trials.

In half of the test trials, the attended stream had the syllable-sequence probabilistic structure implicitly learned by subjects during exposure (i.e., the attended stream consisted only of words from the artificial language to which the subject had been exposed; Predictable-Attended condition; note that consecutive words never repeated). In the remaining trials, the attended stream was a semi-structured sequence of the same syllables (Control-Attended condition). This semi-structured sequence consisted of an equal mix of “part-words” (triplets in which the first two syllables were from a word in the artificial language, and third syllable was unpredicted) and “non-words” (triplets in which the syllable-transition probabilities were completely different from those in the artificial language, with the constraint that the ith syllable of each non-word was always the ith syllable of a language word, for i = 1 to 3; see Figure 1). Part-words and non-words were randomly interleaved within each Control-Attended trial. Further, contextual condition (Predictable-Attended vs. Control-Attended) was randomly interleaved across trials. The attended and ignored streams were staggered in time by a duration of half a syllable, with the attended stream leading in half of the trials and the ignored stream leading in the remaining; attended-lead and attended-lag conditions were randomly interleaved across trials.

The target-stream-in-quiet test task was identical to the target-stream-in-competition task except that the target stream was presented in quiet (without a competing stream).

Each of the two test tasks had 48 trials; each trial lasted ∼27 sec and presented 20 triplets per stream. Thus, the total test duration was ∼30 mins per test task.

For in-lab participants, 32-channel EEG was measured simultaneously with behavior during both exposure and testing. Also, during both exposure and testing, subjects were instructed to stay still while silently counting target syllables in each trial and to only respond at the end of the trial; this procedure mitigates EEG motor artifacts.

#### 4.3.4. Paradigm 2 testing

Paradigm 2 was similar to paradigm 1, differing primarily in the conditions presented in the testing portion of the study. Specifically, paradigm 2 used three contextual conditions during testing: Predictable-Masker, Random-Both, and Predictable-Attended. These three conditions were presented an equal number of times and randomly interleaved across 36 test trials, yielding a total test duration of ∼22 mins. All trials tested syllables in competition (not in quiet) and each subject performed all three contextual conditions in a fully within-subjects design.

In the Predictable-Masker condition, the ignored stream had the syllable-sequence probabilistic structure implicitly learned by subjects during exposure while the attended stream consisted of a random sequence of the same syllables. In the Random-Both condition, both the attended and ignored streams consisted of a random sequence of the same nine syllables. In the Predictable-Attended condition, the attended stream followed the implicitly learned syllable-sequence probabilistic structure while the ignored stream consisted of a random sequence. Consecutive language words never repeated in the predictable stream.

As before, the attended and masker streams were spoken by different talkers. The same two voices that were used during exposure were also used during testing. Thus, in the first experiment of paradigm 2, the first female talker and the male talker were used; in the second experiment, the two female talkers were used. In each experiment, both voices were used for the attended stream, with the attended voice randomly interleaved across trials. Note that while source segregation was expected to be relatively easy in the first experiment, it was expected to be harder in the second experiment because the target and masker voices were more similar in the second experiment.

#### 4.3.5. Online screening

In the online experiments, subjects were instructed to perform the behavioral tasks using their personal computers and headphones/earphones. Headphone checks were performed at the beginning of each online experiment using a paradigm validated by Mok et al. (2024) that we have used in prior studies (Viswanathan et al., 2022; Viswanathan, Heinz, et al., 2024; Viswanathan, Shinn-Cunningham, et al., 2021). In this paradigm, subjects first performed a task to ensure that they were using headphones (Woods et al., 2017). Subjects then performed a second task to test whether headphones/earphones presented signals to both ears. This task was a three-interval three-alternatives-forced-choice task where the target interval contained white noise with interaural correlation fluctuating at 20 Hz, while the dummy intervals contained white noise with a constant interaural correlation. Subjects were asked to detect the interval with the most flutter or fluctuation. Only those subjects who scored greater than 65% in each of the two headphone-check tasks were allowed to proceed to the rest of the experiment.

Subjects performed a level-setting task before the headphone checks and also before the main experimental tasks. In the level-setting task, subjects were asked to make sure that they were in a quiet room and wearing headphones or earphones, and not to use computer speakers. They were then asked to set their computer volume to 10%–20% of the full volume, after which they were played a level-setting stimulus. The level-setting stimulus was a speech-in-babble stimulus if the level setting was performed prior to headphone checks. If the level setting was performed before the main experimental tasks (i.e., the exposure and test tasks), then a level-setting stimulus comprising a random sequence of the same nine syllables used in the main tasks was used, with the talker voice alternating between the two voices used in the main tasks every 15 syllables. Subjects were asked to adjust their volume up to a comfortable, but not too loud level, and were instructed not to change the volume setting for the remainder of the experiment (to avoid sounds becoming too loud or soft). The root-mean-square value of the target stream was matched between the level-setting stimuli and the stimuli used in the tasks following the level-setting task.

### 4.4. Stimulus delivery and EEG recording

A personal desktop computer controlled all aspects of the in-lab experiment, including triggering sound delivery and storing data. A Fireface UFX+ Audio Interface (RME Audio) presented audio diotically through insert earphones (ER-3A; Etymotic) coupled to foam tips. The target and distractor streams were each presented at 71 dB SPL. All audio stimuli were sampled at 44100 Hz. EEG signals were recorded using a Biosemi ActiveTwo system and sampled at 4096 Hz. EEG recordings were done with 32 cephalic electrodes and two additional earlobe electrodes.

### 4.5. Data preprocessing

Subjects who performed at or below theoretical chance levels (p<0.01; Supplementary Figure S5) either during the exposure task or during testing were excluded from the study; six of the in-lab subjects and 11 of the Prolific.com subjects (one from paradigm 1, three from the first experiment of paradigm 2, and seven from the second experiment of paradigm 2) were excluded for this reason. Additionally, in-lab behavioral data from two subjects and EEG from these two subjects and an additional third subject were excluded from the study due to technical issues that arose during data collection. Thus, among the in-lab subjects, behavioral data from 50 (of which 30 performed target stream in competition during testing, and 20 performed target stream in quiet) and EEG data from 49 (of which 29 performed target stream in competition during testing, and the rest target stream in quiet) were included in the study. Among the online subjects, only 48 subjects for paradigm 1, 37 subjects for the first experiment of paradigm 2, and 33 for the second experiment of paradigm 2 were included in the study.

EEG signals were re-referenced to the average of the two earlobe reference electrodes. Signal space projection was used to construct spatial filters to remove eye blink and saccade artifacts (Uusitalo & Ilmoniemi, 1997). Following line noise removal, the broadband EEG was bandpass filtered: 0.1–40 Hz for event-related potential (ERP) calculations, 7–15 Hz for alpha power calculation, 13–30 Hz for beta power calculation, and 0–6 Hz for intertrial phase-locking value (ITPLV) computation. Data were then epoched into non-overlapping time chunks time-locked to the onsets of either syllables, triplets, or a contiguous group of triplets in either stream, then baseline corrected by subtracting the mean in a baseline period (-0.2–0 s relative to the start of the epoch for ERP and spectrum calculations; -0.05–0 s relative to the start of the epoch for ITPLV computation). Zero-group-delay non-causal filtering was used throughout. Epochs with peak-to-peak signal amplitude exceeding 400 microvolts were rejected.

### 4.6. Behavioral and EEG data analysis

Percent trials correct was computed as the percentage of trials in which the reported number of target syllables matched the number presented. Size of error (SOE) was computed as the square root of the mean squared error between the number of targets reported and presented across incorrect trials, and expressed as a percentage relative to the average correct answer in those trials (Equation 1).

Let *P*_*k*_ be the number of target syllables presented in an incorrect trial *k, R*_*k*_ be the number of target syllables reported in trial *k*, and *N* be the number of incorrect trials (i.e., trials where the number of target syllables reported don’t match the number presented). Then,

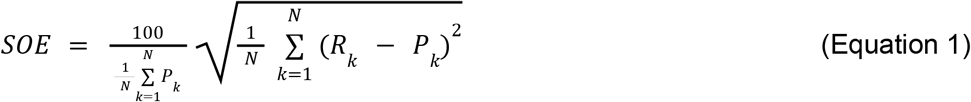

The parietal P300 EEG ERP was computed by averaging EEG data across the parietal channels Pz, CP1, CP2, P3, and P4 and across 73 epochs (which were time locked to the onset of target syllables) per subject and condition. A similar approach was used to calculate fronto-central ERPs (e.g., N100) but using channels Fz, Cz, FC1, and FC2 instead. ERPs were smoothed across time using a third-order-polynomial Savitzky-Golay filter (Schafer, 2011) with a 250-ms window length.

Induced alpha and beta power were calculated as in our prior work (Viswanathan et al., 2023) by computing the EEG spectrogram in the alpha (7–15 Hz) and beta (13–30 Hz) bands, respectively, using 73 epochs (time locked to the onset of triplets in the attended stream) for each subject and condition. A Slepian-tapered complex exponential wavelet, which minimizes spectral leakage (Slepian, 1978; Thomson, 1982), was used. Five cycles were used to estimate each time-frequency bin with a time-full-bandwidth product of 2. The logarithm of the computed spectrogram was taken to ensure response data are normally distributed (Thomson, 1991).

ITPLV (Tallon-Baudry et al., 1996) was computed using an EEG epoch duration of ten triplets, with each epoch starting at the onset of every tenth triplet in the attended stream. The epochs were detrended by removing the mean across time, then the Fourier transform of the result was computed using a Slepian window. Finally, ITPLV was calculated as the degree of phase synchrony of the EEG responses at different frequencies across 48 epochs per subject and condition, as in Equation 2, and then averaged over all 32 channels.

Let *Xi*(*f*) be the Fourier transform of the EEG in epoch *i* at frequency *f*, and let *N* be the total number of epochs. Then,

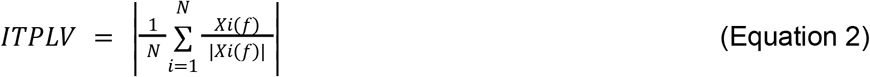

The mean (i.e., expected value) of the theoretical null distribution for ITPLV is 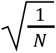 (Bharadwaj & Shinn-Cunningham, 2014; Bokil et al., 2007). Thus, in order to be able to compare ITPLV across conditions, we used the same number of epochs for the ITPLV computation across conditions.

### 4.7. P300 latency computation

Figure 8 illustrates our approach for computing the latency of target-syllable-evoked P300 responses, which is based on prior work (Gorga et al., 2003; Viswanathan, Rupp, et al., 2024). The approach estimates the time at which the neural response begins to deviate from baseline, while at the same time being robust to baseline noise. P300 latency was computed as the time point of initial deviation of the parietal ERP (averaged over channels Pz, CP1, CP2, P3, and P4) 180 ms post target-syllable onset, leveraging a leave-one-out jackknifing procedure (Tukey, 1958) to obtain stable estimates for the mean and standard error of the latency estimates across subjects. Specifically, the mean and 95% confidence interval (CI) in a baseline period beginning at the onset of each triplet and ending 180 ms post the onset of the third/target syllable was first computed. Then the earliest time point (T1) at which the average parietal ERP 1) exceeds the baseline 95% CI (baseline 95% CI = baseline mean + 1.96 times the standard error of the baseline mean; note that 1.96 corresponds to the 95% CI for the distribution under the null hypothesis that there is no P300 response) and 2) remains above baseline for at least 30 ms was calculated. Finally, a straight line (L1) was fitted with data points -5 to 10 ms around T1, and the time at which L1 crosses the baseline mean was taken to be the latency. The combination of our pre-processing approach (especially, zero-group-delay filtering of the raw EEG) and latency computation procedure renders our latency estimates invariant to any response smearing in time introduced by the filtering process.

**Figure 8.**
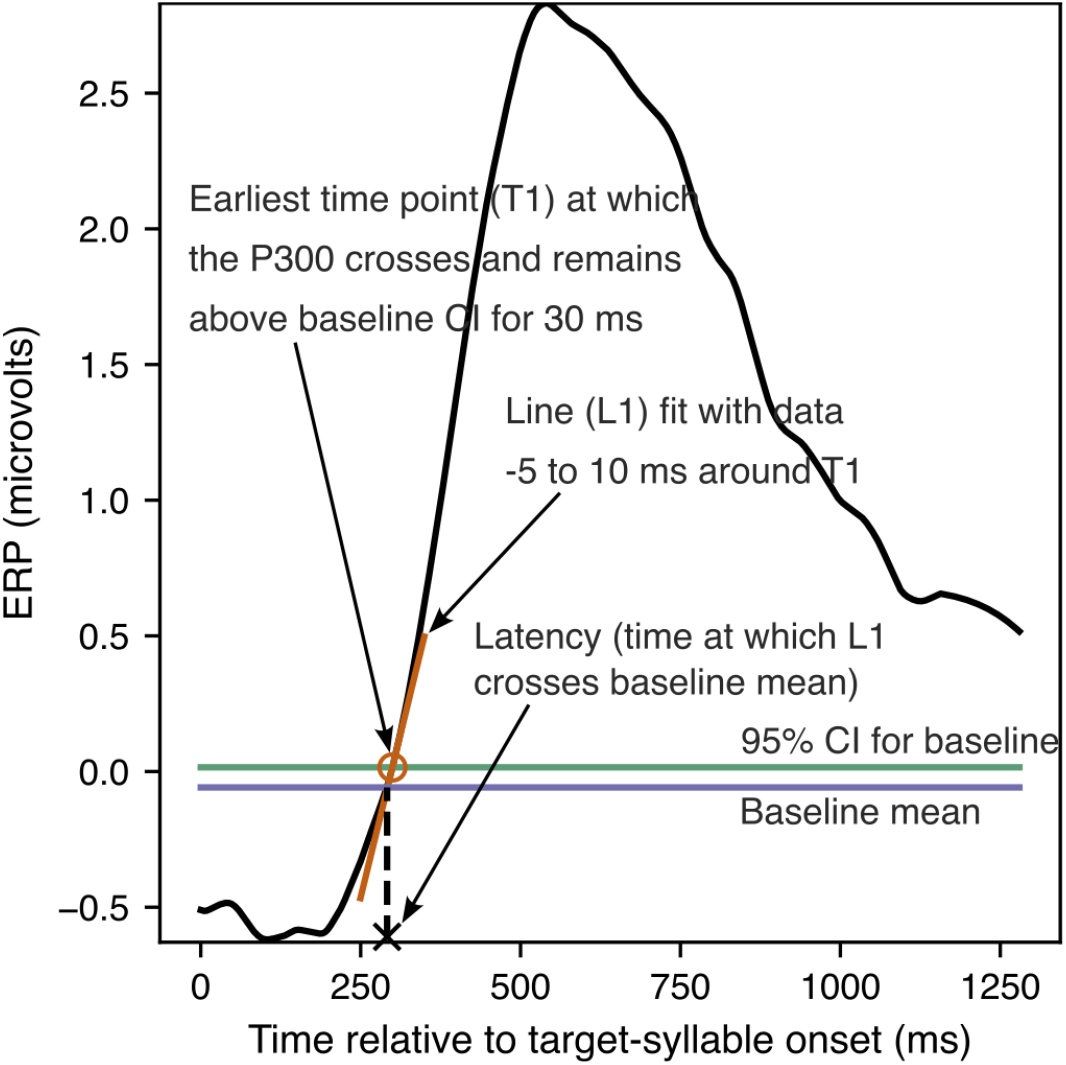
Schematic of P300 latency computation. The earliest time point (T1; orange circle) 180 ms following target-syllable onset at which the average parietal ERP crosses the baseline (time period between triplet onset and 180 ms post target-syllable onset) 95% confidence interval (CI; green line) and remains above the CI for at least 30 ms was computed. A straight line (L1; orange line) was fit with the set of data points -5 ms before to 10 ms after T1. The time at which L1 crosses the baseline mean (purple line) was taken to be the P300 ERP latency (black cross).

### 4.8. Statistical analysis

Nonparametric permutation testing (Nichols & Holmes, 2002) was used for all behavioral data to account for any ceiling effects that may cause the data to have a non-Gaussian distribution. To test whether target syllables are easier to detect in the Predictable-Attended than the Control-Attended contextual condition (paradigm 1) and whether they are easier to detect in the Predictable-Attended and Predictable-Masker contextual conditions compared to the Random-Both condition (paradigm 2), we computed the prediction benefit on percent trials correct and SOE as the within-subject percent change in percent trials correct and SOE, respectively, between the different contextual conditions under comparison. To generate a single realization from the distribution under the null hypothesis of zero prediction benefit, the contextual-condition labels (e.g., Predictable-Attended and Control-Attended) were swapped randomly for each subject, then the average prediction benefit across subjects was computed with the new condition labels. This procedure was repeated with 100,000 distinct randomizations to generate each full null distribution of zero prediction benefit. The resulting null distributions were used to assign p-values to the observed average prediction benefits obtained with the correctly labeled data.

A between-subjects nonparametric permutation testing procedure was used to test whether the prediction benefit for target-syllable detection is greater in competition than in quiet in experiment 1 of paradigm 1 since different subjects performed the two masking conditions (in competition vs. in quiet) during testing. To generate a single realization from the distribution under the null hypothesis of zero difference in prediction benefit across masking conditions, each subject was randomly assigned a masking-condition label such that the total number of subjects per masking condition was the same as in the original data. Then the average prediction benefit was computed across subjects for each masking condition and its difference taken across masking conditions. This procedure was repeated with 100,000 distinct randomizations to generate the full null distribution of zero difference in prediction benefit across masking conditions. The generated null distribution was used to assign a p-value to the observed average prediction-benefit difference obtained with the correct masking-condition labels. A similar approach was used to test for an interaction between contextual and masking conditions in the online replication experiment (experiment 2) of paradigm 1, but leveraging the fully within-subjects online experiment design. Specifically, to generate a single realization from the null distribution, a random sign was assigned to the within-subject difference in the prediction benefit between masking conditions, then the results were averaged across subjects. As before, this procedure was repeated with 100,000 distinct randomizations to generate the full null distribution of zero difference in prediction benefit across masking conditions, and the generated null distribution was used to assign a p-value to the average prediction-benefit difference obtained with the correct masking-condition labels.

Similarly, within-subject permutation tests were used to determine whether the difference in the number of targets syllables presented and reported on average (across trials) varies between contextual conditions (e.g., comparing Control-Attended with Predictable-Attended in paradigm 1, and comparing Predictable-Masker and Random-Both with Predictable-Attended in paradigm 2).

A paired t-test was used to test whether the P300 ERP latency is smaller for predicted (Predictable-Attended condition) targets compared to unpredicted (Control-Attended condition) targets within any given masking condition (in competition, or in quiet). The Welch two-sample t-test (Welch, 1947) was used to test whether the P300 ERP latency is smaller for predicted targets in competition compared to unpredicted targets in quiet.

To test whether ITPLV is greater for the Predictable-Attended condition than the Control-Attended condition at different frequencies (specifically, the triplet rate, the first harmonic of the triplet rate, and the syllabic rate), the ITPLV was first averaged over all channels and over frequency bins 0.2 Hz (the approximate frequency resolution of the ITPLV calculation) around the frequencies of interest. Then, the percent change in the ITPLV between Predictable-Attended and Control-Attended conditions was computed and a one-sample t-test was used to test whether this result is significantly greater than zero.

A one-sample t-test was used to analyze whether the ERP amplitudes between the onsets of the second and third syllables differ significantly between the cases when the target is predicted by the preceding two syllables compared to when the first but not second syllable predicts the target. For each subject, the channel-averaged ERP difference between these two conditions was computed and averaged across time between the onsets of the second and third syllables. We tested whether the mean of the resulting across-subject distribution is significantly different from zero.

## Acknowledgements

This study was supported by the Office of Naval Research [MURI grant number N00014-23-1-2065 (to B.G.S.-C.)] and the National Institutes of Health [grant R01HD105313 (to J.R.S.)].

## 4.9. Data and software availability

Stimuli were created using MATLAB (The MathWorks, Inc., Natick, MA) and Praat (Boersma, 2007). Stimulus presentation was controlled using custom MATLAB routines. EEG preprocessing and calculation of power spectral density and ITPLV were performed using the open-source software tools MNE-PYTHON (Gramfort et al., 2014) and SNAPsoftware (Bharadwaj, 2021). All further data analyses were performed using custom software in PYTHON (Python Software Foundation, Wilmington, DE) and MATLAB. Statistical analyses were performed using R (R Core Team; www.R-project.org). Visualizations used color-blind-friendly Colorbrewer colormap palettes (Brewer et al., 2003). Our custom code is available publicly at https://github.com/vibhaviswana/Viswanathan2026_predictionFromStatisticalLearningAidsSceneAnalysis.

