## Supplementary Figures for "Prediction from Statistical Learning Aids Auditory Scene Analysis"

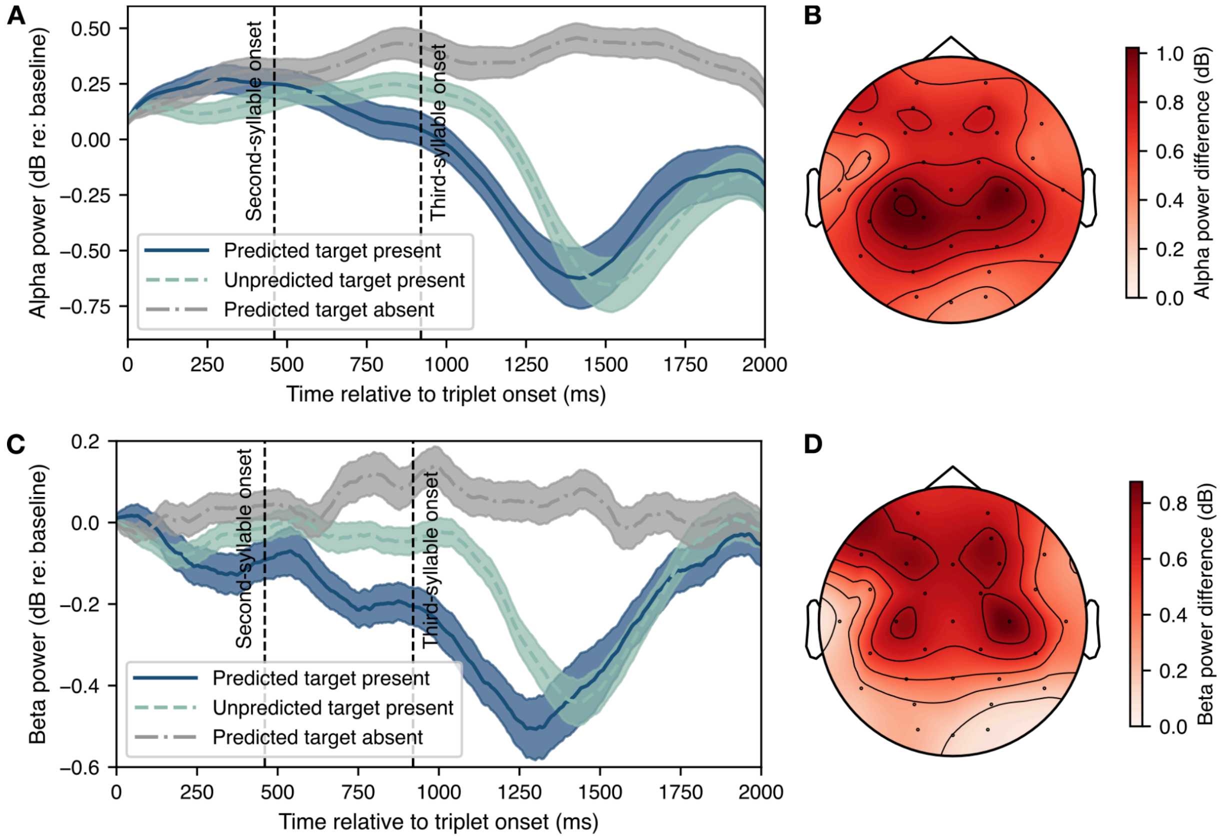


**Figure S1.** Induced alpha and beta power (paradigm 1 test-stage EEG data pooled over masking conditions). Panels A and C show alpha and beta power time courses (in dB relative to the mean in a baseline period -50 to 0 ms post triplet onset), respectively, for attended-stream triplets in the Predictable-Attended and Control-Attended/Unpredictable conditions and for the cases when the third syllable is the target and when it is not the target (mean and standard error across subjects). Data shown are averaged over epochs, either parieto-occipital (A9, A10, A11, A12, A13, A14, A15, A16, A17, A18, A19, A20, A21, and A22; for alpha) or frontal (A1, A2, A3, A4, A5, A6, A25, A26, A27, A28, A29, A30, and A31; for beta) channels, and band-specific frequency bins. Time zero corresponds to the onset of the first syllable; the onsets of the second and third syllables are marked. Panels B and D show scalp topomaps of the alpha and beta power differences, respectively, between the cases when the target syllable is absent and when it is present (in both cases, the target is predicted by the preceding syllables). Data shown are averaged across epochs, frequency bins, subjects, and time between 0 and 1 s post third-syllable onset.

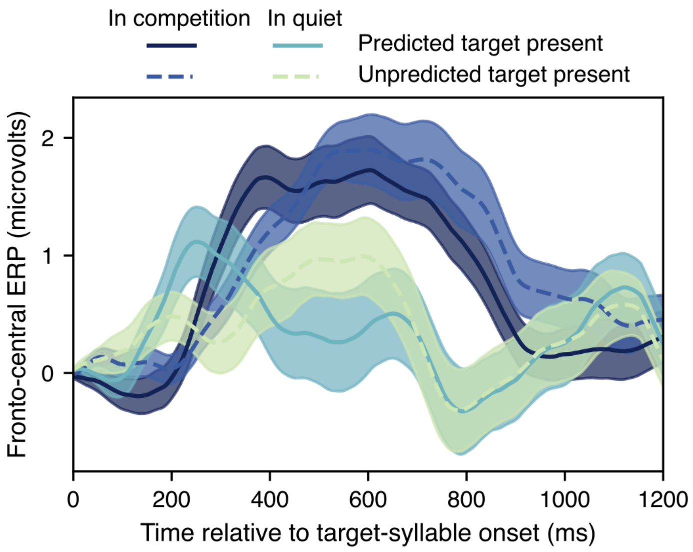


**Figure S2.** Average ERP timecourse in fronto-central electrodes (Fz, Cz, FC1, FC2, CP1, and CP2) in the different contextual (Predictable-Attended or Control-Attended/Unpredictable) and masking (in-competition or in-quiet) conditions (paradigm 1 test-stage EEG data; mean and standard error across subjects). The target syllable is present in all cases shown.

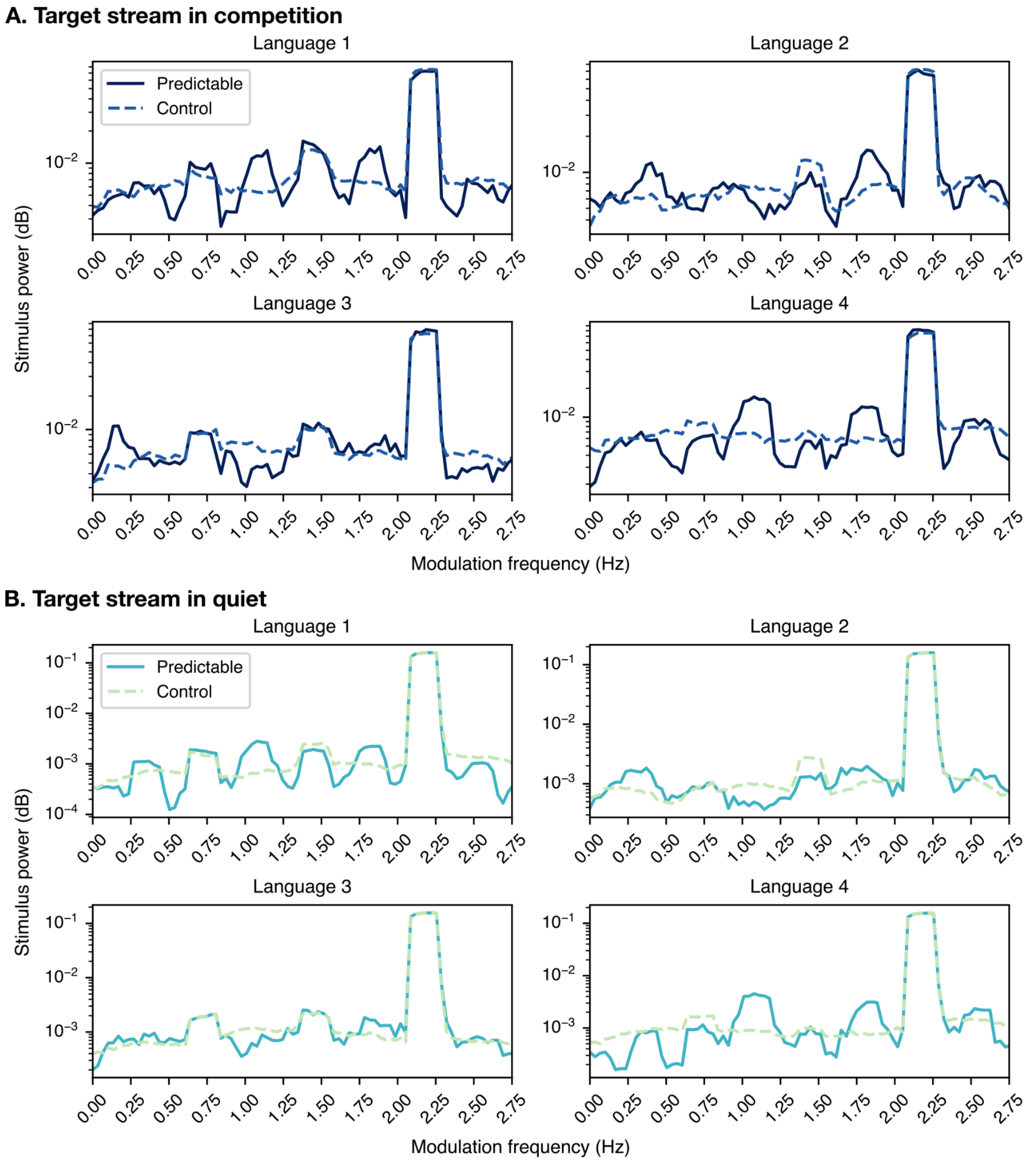


**Figure S3.** Modulation spectra of the target stream in competition (Panel A) and in quiet (Panel B) stimuli for each of the four artificial languages from paradigm 1. Data are shown separately for the Predictable-Attended and Control-Attended conditions. Peaks in the modulation spectra are present at the syllabic rate (2.17 Hz), triplet rate (0.72 Hz), and their harmonics. Peaks at the two-syllable (spondee) rate (1.1 Hz), the 6-syllable (2-word) rate (0.36 Hz), and their harmonics are also seen in some of the artificial languages in the Predictable-Attended condition.

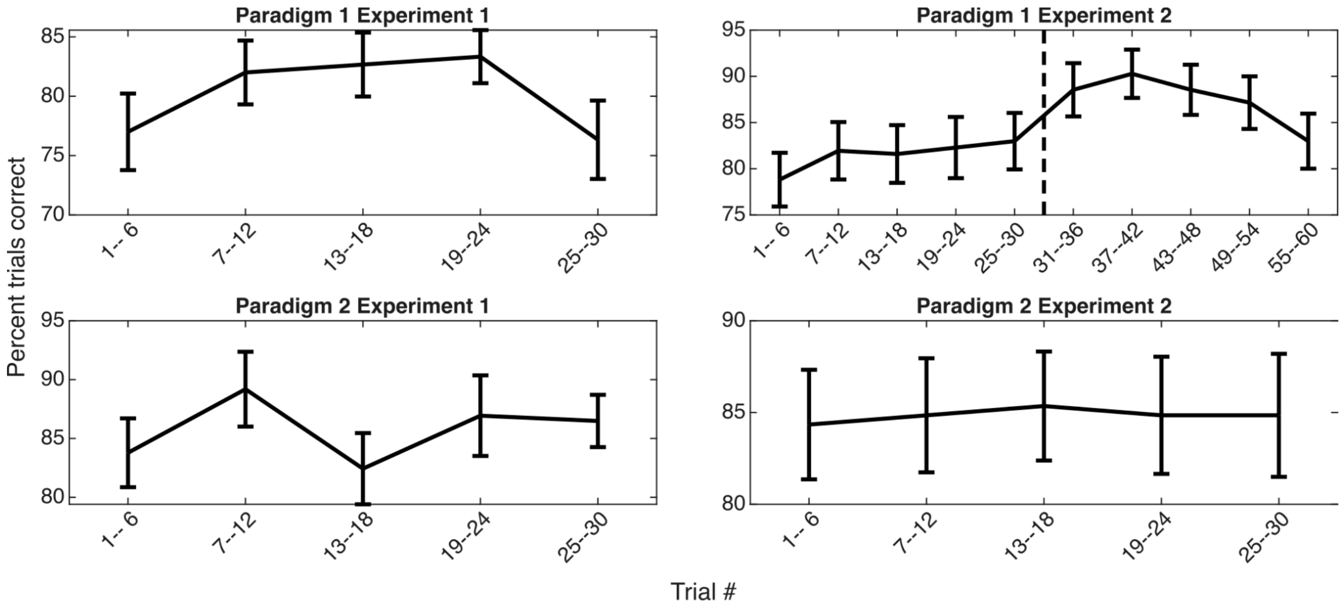


**Figure S4.** Exposure behavioral performance (percent trials correct) as a function of exposure duration (mean and standard error across subjects). Data are plotted for contiguous exposure trial chunks, with each trial chunk consisting of 6 contiguous trials. Note that experiment 2 of paradigm 1 had two exposure stages with a break in between the stages (indicated by a vertical dashed line); this break had an average duration of 7 days and most subjects had a break close to 5 or 11 days, while 3 out of the 48 subjects did the second exposure on the same day as the first exposure.

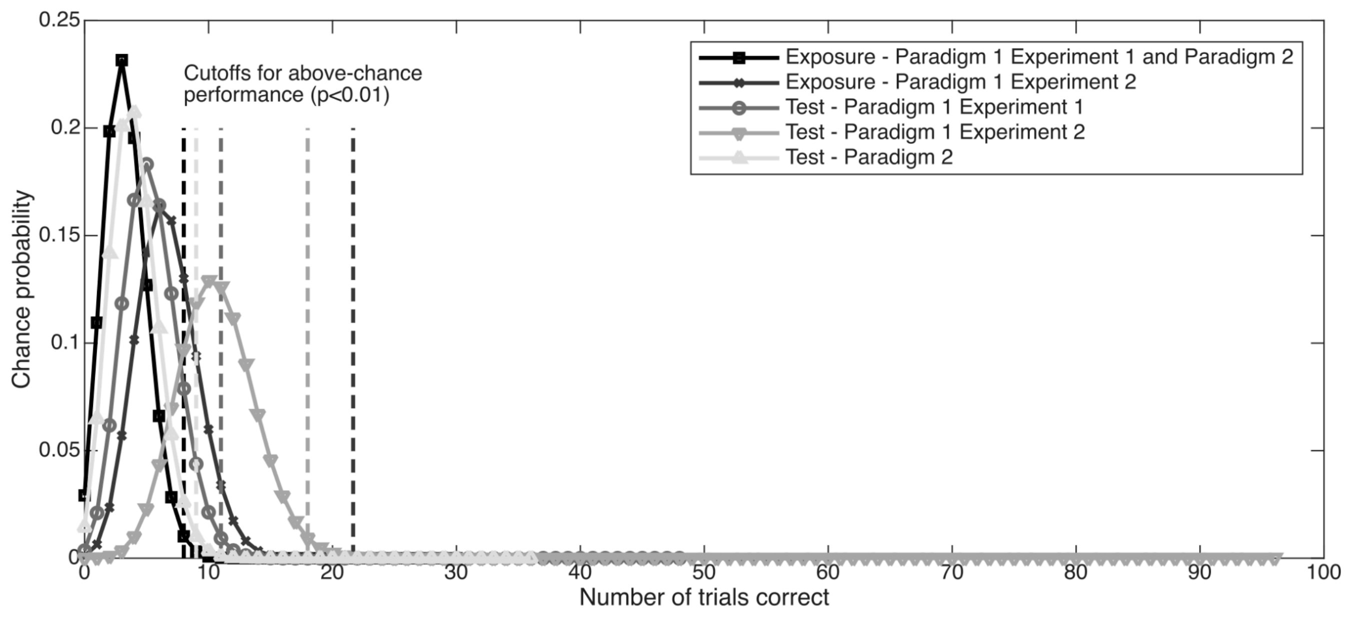


**Figure S5.** Theoretical chance probability for number of trials correct for the exposure and test tasks of each paradigm. The chance probability is a Binomial distribution parameterized by the total number of trials in the task and the probability that any given trial is correct. The total number of trials equals 30 for the exposure task in paradigm 1 experiment 1 and in paradigm 2, 60 across the two exposure tasks of paradigm 1 experiment 2, 48 for the test task in paradigm 1 experiment 1, 96 across the two test tasks in paradigm 1 experiment 2, and 36 for the test task in paradigm 2. The probability that any given trial is correct is the reciprocal of the number of choices for the number of targets presented per trial (the choices were 0 through 8, and so the total number of choices equals 9).
